# Dogs and cats are less susceptible to the omicron variant of concern of SARS-CoV-2 – a field study

**DOI:** 10.1101/2023.02.07.527419

**Authors:** Constantin Klein, Anna Michelitsch, Valerie Allendorf, Franz Josef Conraths, Martin Beer, Nicolai Denzin, Kerstin Wernike

## Abstract

Severe acute respiratory syndrome coronavirus 2 (SARS-CoV-2) caused a pandemic of unprecedented extent. Beside humans, a number of animal species can be infected, however, in some species differing susceptibilities were observed depending on the virus variant. Here, we serologically investigated cats and dogs living in households with human COVID-19 patients. The study was conducted during the transition period from delta as the dominating variant of concern (VOC) to omicron (BA.1/BA.2) to investigate the frequency of virus transmission of both VOCs from infected owners to their pets. The animal sera were tested by surrogate virus neutralization tests (sVNT) using either the original receptor-binding domain (RBD), enabling the detection of antibodies against the delta variant, or an omicron-specific RBD. Of the 290 canine samples, 20 tested positive by sVNT, but there were marked differences between the sampling time, and, related thereto, the virus variants, the dogs had contact to. While in November 2021 infected owners led to 50% seropositive dogs (18/36), only 0.8% (2/254) of animals with household contacts to SARS-CoV-2 between December 2021 and April 2022 tested positive. In all cases, the positive reaction was recorded against the original RBD. For cats, a similar pattern was seen, as in November 2021 38.1% (16/42) tested positive and between December 2021 and March 2022 only 5.0% (10/199). The markedly reduced ratio of seropositive animals during the period of omicron circulation suggests a considerably lower susceptibility of dogs and cats to this VOC.

To examine the effect of BA.2, BA.4 and BA.5 omicron subvariants, sera taken in the second and third quarter of 2022 from randomly selected cats were investigated. 2.3% (11/372) tested seropositive and all of them showed a stronger reaction against the original RBD, further supporting the assumption of a lower susceptibility of companion animals to the omicron VOC.

## Introduction

Severe acute respiratory syndrome coronavirus 2 (SARS-CoV-2), a betacoronavirus of the subgenus *Sarbecovirus*, is the causative agent of the human disease COVID-19 (= coronavirus disease 2019), which was reported for the first time in late 2019 in Wuhan, China (Zhu et al., 2020). Thereafter, the novel pathogen very rapidly spread globally, driven by direct human-to-human virus transmission via aerosolized particles, and led to millions of human deaths worldwide (Dong, Du, & Gardner, 2020; WHO, 2020). Since the first detection of the virus, SARS-CoV-2 has evolved, leading to the emergence of numerous variants, some of which represent so-called variants of interest (VOI) or variants of concern (VOC) that are continuously monitored by e.g. the World Health Organization (WHO, 2022b). VOCs could spread more easily, escape the host’s immune response, cause more severe disease in humans, change the clinical presentation, decrease the effectiveness of vaccines, treatments or diagnostic tools, or display an altered host range (WHO, 2022a).

Beside humans, several animal species can be infected with SARS-CoV-2, among them non-human primates, felines, canines, mustelids, some ruminant species and several rodents (OIE, 2021), but their susceptibility might vary depending on the virus variant. As an example, house mice (*Mus musculus*) were excluded as amplifying host for the wild-type virus by experimental infection (Bosco-Lauth et al., 2021), but they are susceptible to some VOCs, in particular the beta (B.1.351) and gamma (P1) variants (Montagutelli et al., 2021; Tarrés-Freixas et al., 2022). Ferrets (*Mustela putorius*) are susceptible to wildtype SARS-CoV-2 and some of the VOCs, particularly alpha (B.1.1.7) and delta (B.1.617.2) (Barut et al., 2022; Pulit-Penaloza et al., 2022; Schlottau et al., 2020; Shi et al., 2020; Ulrich et al., 2021). However, a BA.1 strain of the omicron (B.1.1.529) VOC failed completely to induce productive infections in these animals (Barut et al., 2022).

Among the animal species in principle susceptible to SARS-CoV-2 those are of particular concern that are frequently in close contact to humans, such as companion animals like cats (*Felis catus*) and dogs (*Canis lupus*). Therefore, pets were included in surveillance studies early in the course of the human pandemic, and, indeed, anthropo-zoonotic SARS-CoV-2 transmissions from infected owners to their cats or dogs were noticed and included multiple VOCs (Carneiro et al., 2022; Hamer et al., 2022; Jairak, Chamsai, et al., 2022; Miró et al., 2021). In various seroprevalence studies, varying proportions of seropositive animals were found, which depended on the location and especially the study period. During the first months of the pandemic, relatively small proportions of seropositive animals (< 5 %) were found in randomly selected samples, while higher seroprevalences were observed in pets as the pandemic progressed and the number of cases in humans increased sharply worldwide (Dileepan et al., 2021; Ito et al., 2021; Jairak, Charoenkul, et al., 2022; Kaczorek-Łukowska et al., 2022; Michelitsch, Hoffmann, Wernike, & Beer, 2020; Patterson et al., 2020). Particularly high seroprevalences were recorded when cats and dogs from households with human COVID-19 patients were sampled (Fernández-Bastit et al., 2022; Kannekens-Jager et al., 2022; Meisner et al., 2022).

Although a few infections of companion animals with the omicron VOC were published (Piewbang et al., 2022; Sánchez-Morales, Sánchez-Vizcaíno, Pérez-Sancho, Domínguez, & Barroso-Arévalo, 2022), often reporting low viral loads, there are considerably fewer reports than during the periods at which the earlier VOCs were circulating. So far, it is not known whether cats and dogs are less susceptible to omicron, especially as data from experimental infection studies are not yet available, or whether only frequency of testing and reporting has decreased. Therefore, we investigated cats and dogs from COVID-19 households during the transition period from delta as the dominating VOC to omicron in the human population, in order to investigate the frequency of virus transmission of both variants from infected owners to their pets.

## Materials and Methods

We aimed to investigate dogs and cats from households with confirmed human COVID-19 cases by serological methods. Households were recruited through information letters distributed by associations of independent veterinarians, social media, online platforms and word of mouth. Serum or plasma samples of the animals were taken between three weeks and three months after the first SARS-CoV-2 detection in a pet owner. The study period started in November 2021, when delta was the dominant variant detected in the human population in Germany, and went through April 2022, thereby covering the period at which omicron BA.1 and BA.2 strains dominated in humans (Robert-Koch-Institut, 2022a) (Figure 1). A total of 241 cats and 290 dogs could be recruited for the study. The number of samples per species and month is given in Figures 2 and 3.

**Figure 1:**
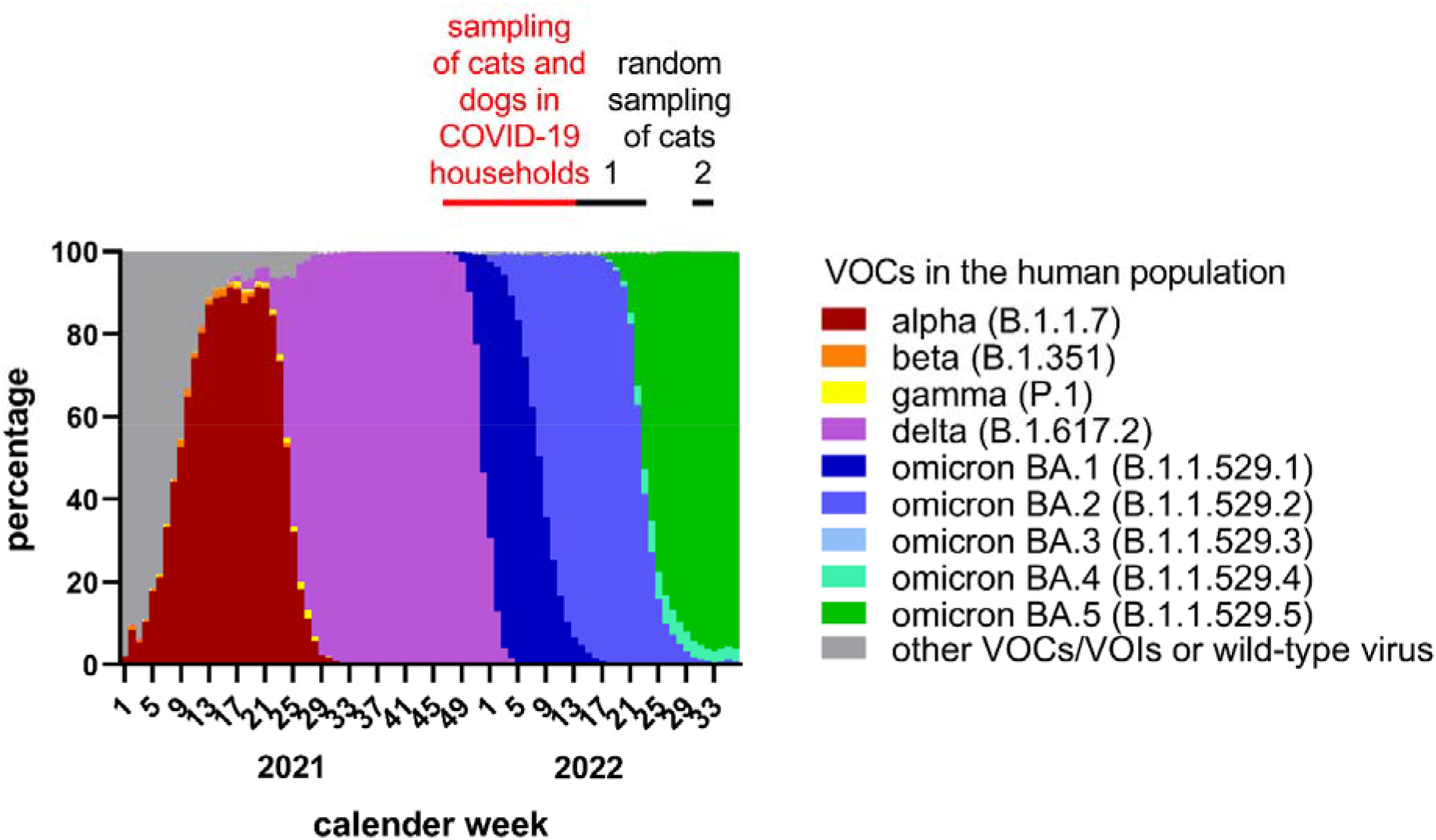
Periods at which samples were taken in the present study and shares of variants of concern (VOCs) detected in human samples in Germany (data retrieved from https://www.rki.de/DE/Content/InfAZ/N/Neuartiges_Coronavirus/Daten/VOC_VOI_Tabelle.html). The sampling period in COVID-19 households is indicated in red and the two time frames at which randomly selected cats were sampled are marked by black bars.

**Figure 2:**
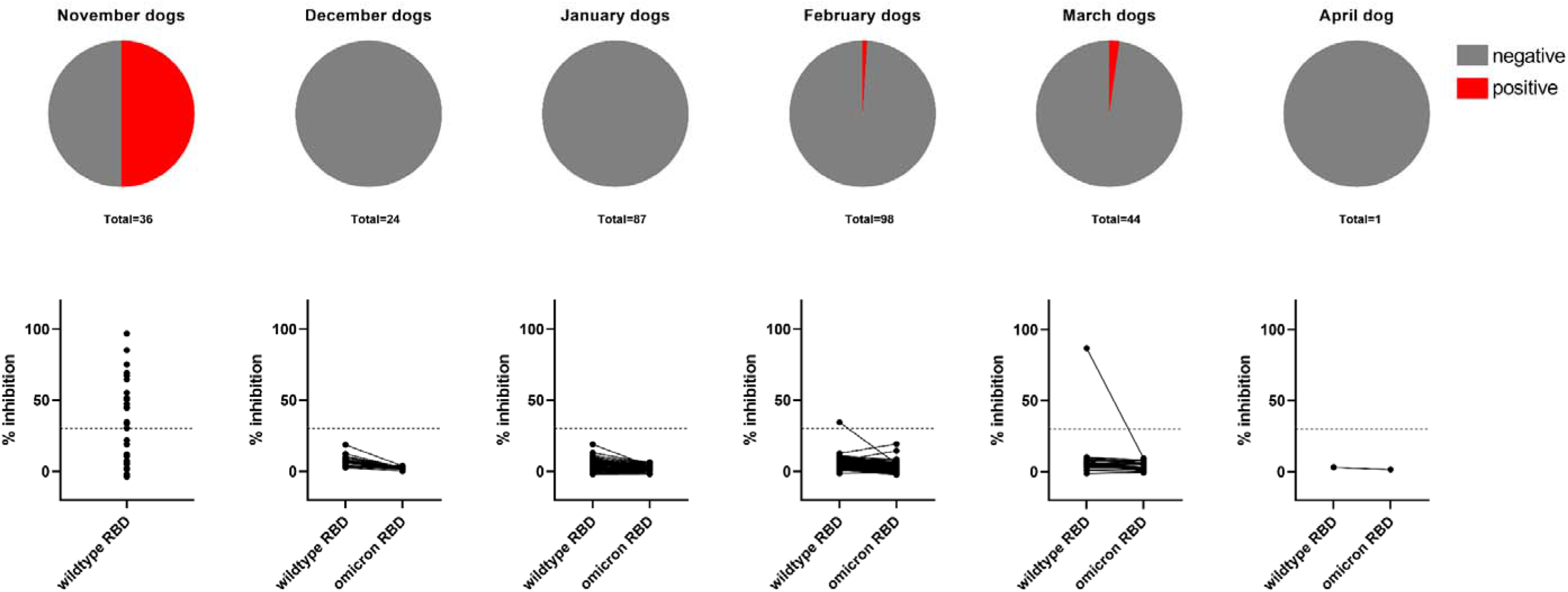
Serological results of dogs kept in COVID-19 households. In the upper panel, the shares of positive (red) and negative (grey) results are given, the animals are sorted into the month in which their owner tested SARS-CoV-2 positive. In the lower panel, the values as measured in the surrogate virus neutralization test are shown individually for each canine sample. In November 2021, before the omicron variant of concern was detected for the first time in the human population of Germany, the canine sera were tested only against the original RBD. From December 2021 onwards, the samples were tested in parallel using the original as well as the omicron RBD and the results of individual samples are connected by a black line. The cut-off is indicated by a horizontal dashed line.

**Figure 3:**
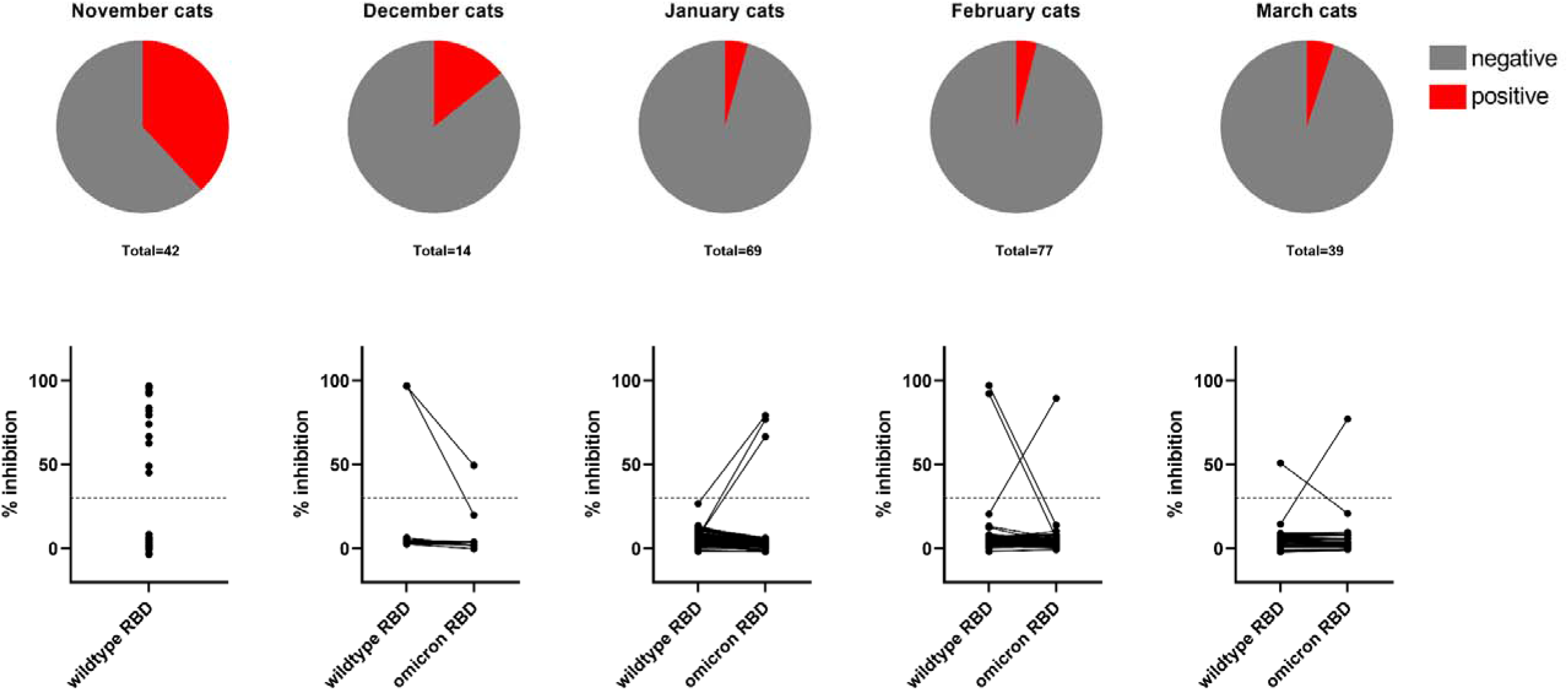
Serological results of cats kept in COVID-19 households. In the upper panel, the shares of positive (red) and negative (grey) results are given, the animals are sorted into the month in which their owner tested SARS-CoV-2 positive. In the lower panel, the values as measured in the surrogate virus neutralization test are shown individually for each feline sample. In November 2021, before the omicron variant of concern was detected for the first time in the human population of Germany, the feline sera were tested only against the original RBD. From December 2021 onwards, the samples were tested in parallel using the original as well as the omicron RBD and the results of individual samples are connected by a black line. The cut-off is indicated by a horizontal dashed line.

To cover the period when omicron BA.4 and BA.5 strains, respectively, were predominantly spreading, randomly selected feline serum or plasma samples taken in Germany between calendar week 11 and 23 of the year 2022 (n=172) or in weeks 31/32 of 2022 (n=200) were serologically tested. The samples were obtained from domestic cats during clinical examination by the attending veterinarian and were sent to a clinical diagnostic laboratory (Laboklin GmbH & Co.KG) for non-SARS-CoV-2-related testing (e.g. hematology testing). Superfluous sample material was kindly provided to the Friedrich-Loeffler-Institut for SARS-CoV-2 serology. The health status of the cats is not known to the authors, the only data given was the postal code of the veterinarian who had taken the sample; the veterinary practices were spread all over Germany. The infection status of the animal owners was likewise unknown, however, given the high prevalence in the human population during the sampling period (Robert-Koch-Institut, 2022b), a certain proportion of cats with contact to infected owners could be expected.

All sera were tested by a commercial, species-independent surrogate virus neutralization test (sVNT) (cPass^™^ SARS-CoV-2 Neutralization Antibody Detection Kit, GenScript, the Netherlands). The test was performed as prescribed by the manufacturer using a cut-off of ≥ 30 % for positivity and < 30 % for negativity. The sVNT in its original composition allows for the detection of antibodies against the wildtype virus and diverse SARS-CoV-2 VOCs including delta, but except omicron. For omicron and its sub-variants, a specific horse radish peroxidase (HRP)-conjugated receptor-binding domain (RBD) protein is provided by the test manufacturer. The suitability to detect and discriminate antibodies against the delta and omicron variants was proven by testing sera obtained from goats experimentally delta-infected (Ulrich et al., unpublished) and from mice infected with an omicron BA.1 strain (kindly provided by the Institute of Virology, Medical Center University of Freiburg, Germany).

Feline and canine samples collected in November 2021, i.e. prior to the first detection of the omicron VOC in the human population of Germany (Robert-Koch-Institut, 2022a) (Figure 1), were tested only by the original composition of the sVNT. Samples collected from December 2021 onwards were tested by the sVNT using the original RBD and, in a parallel approach, using the omicron-specific RBD.

Sera that reacted positive in the sVNT were subsequently tested by an indirect immunofluorescence assay (iIFA) for confirmation. The test was performed as described previously (Michelitsch, Hoffmann, Wernike, & Beer, 2020; Schlottau et al., 2020) with a starting dilution of 1/16 and four log2 dilution steps. As secondary antibody, FITC-labelled anti-cat IgG (dilution 1/600; Sigma–Aldrich, Steinheim, Germany) and anti-dog IgG (1/100; Sigma–Aldrich), respectively, was used.

## Results and Discussion

### Analysis of samples from COVID-19 households suggests a lower susceptibility of dogs and cats to the omicron VOC

Twenty of the 290 dogs that had contact to their SARS-CoV-2 infected owner tested positive by sVNT (6.9%, 95% confidence interval (CI): 4.0% - 9.8%). However, there were marked differences between the months of the study period and, related thereto, between the virus variants the dogs had contact to. While antibodies against SARS-CoV-2 could be detected in 18 of 36 dogs (50.0%, 95% CI: 33.7% - 66.3%) that had contact to an infected owner in November 2021, only 2 of the 254 animals that were sampled from December 2021 through April 2022 tested positive (0.8%, 95% CI: 0.0% −1.9%) (Figure 2). In all cases, the positive reaction was recorded against the original RBD. When using the omicron RBD, every canine sample tested negative (Figure 2). All positive sVNT results could be confirmed by the iIFA as every sample that tested positive in the sVNT also gave a positive result in the iIFA; the titers ranged from 1/32 to >1/128.

For cats, a similar pattern was seen. Overall, 26 of the 241 feline sera (10.8%, 95% CI: 6.9% −14.7%) tested seropositive. Sixteen of the positive reacting sera were collected in households that had COVID-19 patients in November 2021 (16/42; 38.1%, 95% CI: 23.4% - 52.8%). Of the 199 cats that had contact to a SARS-CoV-2 positive owner between December 2021 and March 2022, only 10 scored positive (10/199; 5.0%, 95% CI: 2.0% - 8.1%). From these 10 samples, five showed a stronger reaction against the original RBD and five against the omicron RBD (Figure 3). Again, all positive sVNT results could be confirmed by the iIFA. The titers ranged from 1/64 to >1/128.

To control for the suitability of detecting and discriminating antibodies against the delta and omicron variants, sera obtained from goats experimentally delta-infected and from mice infected with an omicron BA.1 strain were tested. Every control sample reacted in the sVNT as expected, i.e. the goat sera tested positive when using the original RBD, and the mouse samples reacted positive against the omicron RBD (Figure 4).

**Figure 4:**
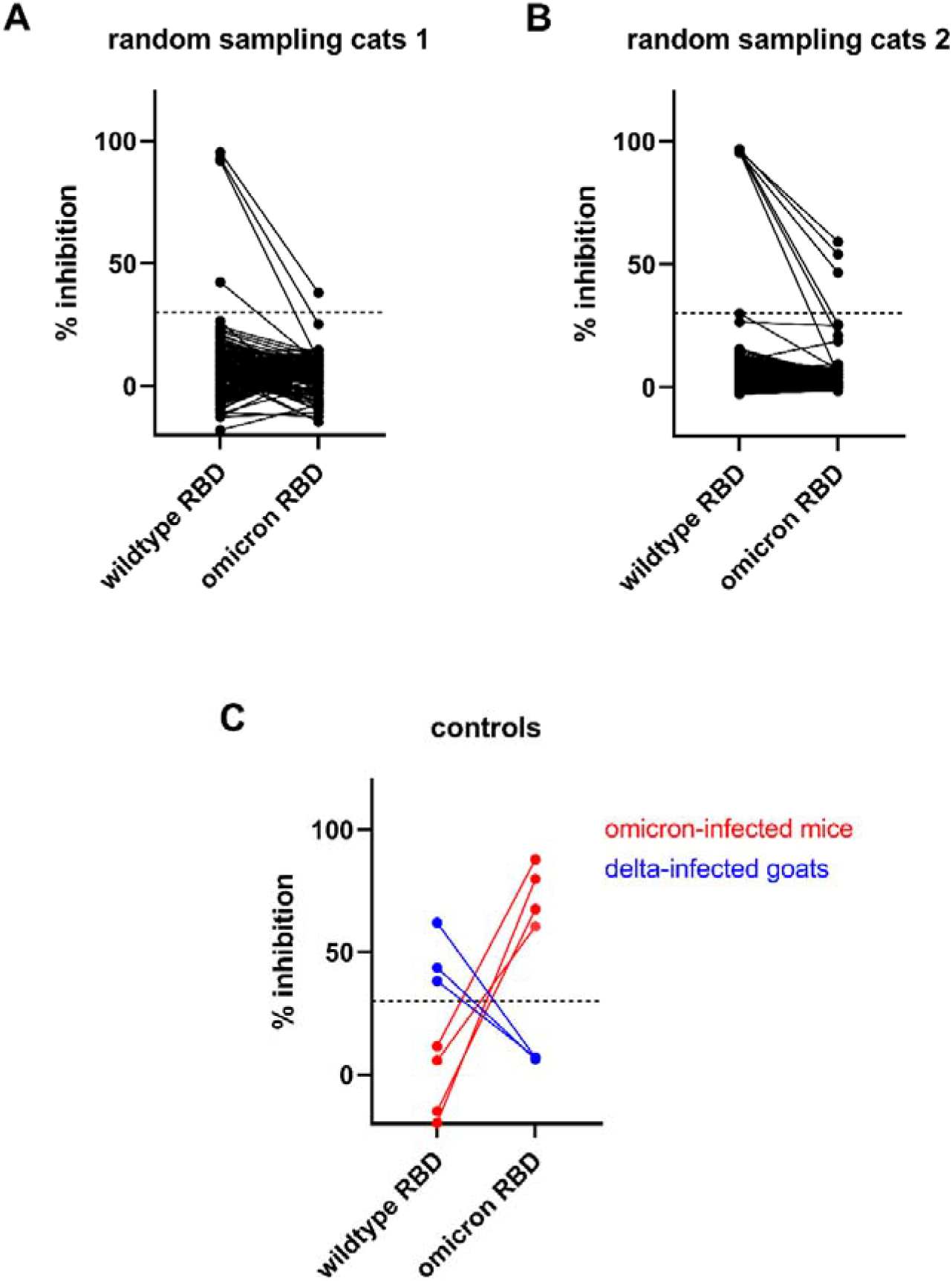
Results of the surrogate virus neutralization test for randomly selected cats sampled in calendar weeks 11 to 23 of 2022 (A) or in calendar weeks 31/32 of 2022 (B) and of the positive control sera (C). All samples were tested against the original as well as the omicron RBD and the results of individual samples are connected by a line. The cut-off of the test is indicated by a horizontal dashed line.

The serological results of cats and dogs, specifically the markedly reduced rate of seropositive animals in the period of omicron circulation in comparison to the delta period, suggest a considerable reduction in the susceptibility of these animal species to the omicron VOC. In November 2021, i.e. when delta represented the dominant VOC in the human population of Germany, a high proportion of cats and dogs was infected by contact to their virus-positive owners. This is in line with previous household studies conducted prior to 2022, where seroprevalences well over 10 % were consistently found, independent of the study area, time and VOC circulating at that time (Fernández-Bastit et al., 2022; Fritz et al., 2021; Goryoka et al., 2021; Meisner et al., 2022). Therefore, cats and dogs appear to be susceptible to the wildtype virus and the earlier VOCs up to delta, which was confirmed for the wild-type virus also by experimental infection (Bosco-Lauth et al., 2020; Shi et al., 2020). In contrast, from December 2021 onwards, when omicron took over the dominance in humans, we detected antibodies against SARS-CoV-2 in only a considerably lower proportion of cats and, among the dogs, not even a single seropositive animal was found. Hence, cats and dogs seem to be much less receptive to omicron, which mirrors the situation in ferrets that belong like cats and dogs to the order Carnivora and that can be infected with the wild-type virus, some of the earlier VOCs, but not with the omicron BA.1 VOC (Barut et al., 2022; Pulit-Penaloza et al., 2022; Schlottau et al., 2020; Shi et al., 2020; Ulrich et al., 2021). This susceptibility pattern in carnivores is in stark contrast to humans, where the secondary attack rate of the omicron variant in households was even higher than that of the delta variant (Jalali et al., 2022).

Ever since the first reporting of omicron in South Africa, its origin was up to debate, with hypotheses about an animal origin and those about continuous evolution in humans being raised (Berkhout & Herrera-Carrillo, 2022; Du, Gao, & Wang, 2022; Wei et al., 2021). The markedly reduced susceptibility or even unresponsiveness of carnivores contradicts the theory of omicron’s origin in an animal reservoir, at least for carnivores, and strongly supports the second hypotheses, i.e. that omicron and its subvariants evolved in and adapted to humans.

### No detection of antibodies against the omicron VOC in randomly sampled cats

Sample collection from cats and dogs from COVID-19 households ended in April 2022. Nevertheless, to also examine the effect of BA.2, BA.4 and BA.5 omicron subvariants on cats, sera of randomly selected animals were tested. For this part of the study, only cats were chosen as this species showed, in contrast to dogs, single seropositive reactions against the omicron RBD in the household study during the time of BA.1/BA.2 circulation. During the first sampling period (calendar weeks 11 to 23 of the year 2022), 172 sera were analyzed and four of them tested positive in the sVNT (2.3%, 95% CI: 0.1% to 4.6%). During the second period (calendar weeks 31/32 of 2022), 200 samples were collected and seven scored positive in the sVNT (3.5%, 95% CI: 1.0% - 6.0%). All sera showed a stronger reaction against the original RBD than against the omicron ortholog (Figure 4) and, again, all positive sVNT results were confirmed by the iIFA. Hence, the results of the random sampling confirmed that of the household study, that is, a lower ratio of seroconversions against the omicron VOC. Overall, several studies demonstrated lower prevalences in companion animals with unknown household status (Fernández-Bastit et al., 2022; Kannekens-Jager et al., 2022), but given the high prevalence of infections with omicron’s subvariants in the human population during the first half of the year 2022 (Robert-Koch-Institut, 2022b), at least single seropositive cats are to be expected. Indeed, antibodies against earlier VOCs could be detected, even though the circulation of non-omicron-VOCs was some time ago and although previous studies demonstrated that serum antibody levels decline in cats to the limit of detection within only a few months (Schulz et al., 2021; Zhang et al., 2020). The latter makes it even more surprising that we could detect antibodies directed against the original RBD, while none of the samples showed a stronger reaction against the omicron RBD. Therefore, we conclude that cats are less susceptible to omicron, presumably with no or only marginal differences between omicron’s subvariants. Nevertheless, dogs and cats should be included in monitoring studies or epidemiological investigations also in the future, especially when new virus variants emerge for which the degree of susceptibility of companion animals is not known.

## Acknowledgements

We thank Bianka Hillmann for excellent technical assistance and LABOKLIN GmbH & Co. KG for providing superfluous sample material from their routine diagnostic submissions. This research was supported by intramural funding of the German Federal Ministry of Food and Agriculture provided to the Friedrich-Loeffler-Institut and received additional funding by the German Federal Ministry of Education and Research (InfectControl 2020 Initiative, project COVMon, grant number 03COV16D).

## Author Contribution

Conceptualization: Martin Beer, Nicolai Denzin and Kerstin Wernike; methodology: Kerstin Wernike; formal analysis: Constantin Klein and Kerstin Wernike; investigation: Constantin Klein and Kerstin Wernike; resources: Anna Michelitsch, Valerie Allendorf, Franz Josef Conraths and Nicolai Denzin; writing–original draft preparation: Kerstin Wernike; writing–review and editing: all authors; visualization: Constantin Klein and Kerstin Wernike; supervision: Kerstin Wernike. All authors have read and agreed to the published version of the article.

## Ethics Statement

The serum samples taken from dogs and cats from COVID-19 households were collected by the responsible veterinarians in the context of diagnostic testing, no permissions were needed to collect the specimens. The randomly selected samples were left-over sera collected in a laboratory for veterinary diagnostics. The samples were taken by the responsible veterinarians in the context of the health monitoring of the respected animal, no permissions were needed to collect the specimens.

## Data Availability Statement

The data that support the findings of this study are available from the corresponding authors upon reasonable request.

